# Mapping single-cell rheology of ascidian embryos in the cleavage stages using AFM

**DOI:** 10.1101/2024.12.01.625795

**Authors:** Takahiro Kotani, Tomohiro Matsuo, Megumi Yokobori, Yosuke Tsuboyama, Yuki Miyata, Yuki Fujii, Kaori Kuribayashi-Shigetomi, Takaharu Okajima

## Abstract

During early embryo development, cell division is highly organized and synchronized. Understanding the mechanical properties of embryonic cells as a material is crucial for elucidating the physical mechanism underlying embryogenesis. Previous atomic force microscopy (AFM) studies on developing embryos revealed that single cells of ascidian embryos in the cleavage stage stiffened and softened during cell division. However, how embryonic cells, as a compliant material, exhibit viscoelastic properties during the cell cycle remains poorly characterized. In this study, we investigated the rheological properties of embryonic cells in the animal hemisphere in the cleavage stages using stress-relaxation AFM and approach-retraction force curve AFM techniques. The AFM measurements revealed that developing single cells followed a power-law rheology observed in single-cell rheology *in vitro*. The embryonic cells increased the modulus (stiffness) and decreased the fluidity (the power-law exponent) towards cell division. We found three rheological states in developing embryos during the cell cycle. The correlation between the cell modulus and the fluidity during the cell cycle was collapsed onto a master curve with a negative correlation, indicating that embryonic cells tightly interacting with the neighboring cells conserved the universality of rheological behavior observed in single cells *in vitro*.

**SIGNIFICANCE:** Understanding the rheological properties of embryonic cells is crucial to elucidate the origin and mechanism of the functional and morphological changes in cells during embryogenesis. AFM-based microrheology revealed that single cells of ascidian embryos in the animal hemisphere in the cleavage stages followed a single power-law rheology, which has no characteristic time scale. Furthermore, the rheological parameters, such as cell stiffness and fluidity, were collapsed onto a master curve with a negative correlation during the cell cycle. These rheological behaviors are similar to those observed in single cells in vitro, indicating that the embryonic cells tightly interacting with each other conserved the intrinsic rheological behaviors of isolated single cells.

## INTRODUCTION

Living cells are organized materials that change their intracellular structures during the cell cycle. During cell division, single cells cause substantial changes in cell morphology and mechanical properties (1,2), which interplay through the remodeling of the cytoskeleton in the cell cortex (3‒5). Atomic force microscopy (AFM) has been widely utilized to measure the mechanical properties of single cells (6,7). Previous AFM studies revealed that the cortical actin filaments of a single cell during the cell cycle can polymerize and depolymerize, leading to time-dependent variations in cell stiffness (5,8,9) and rheological properties (4,10).

In embryogenesis, mechanical cues highly regulate cells (11‒16). Recent studies revealed that the rheological properties of embryos play crucial roles in morphological changes in the developmental processes (17‒21). In the cleavage stage of ascidian embryos, cells undergo rapid and synchronous cell divisions, crucial for determining cell deformation in response to intra- and intercellular forces. Previous AFM studies revealed that the recruitment and release of actin filaments in the cell cortical region in developing embryos leads to cell stiffening and softening, exhibiting a periodic mechanical oscillation during embryogenesis (22,23). However, how the cells as a compliant material change the rheological properties during the cell cycle remains poorly characterized.

In this study, we investigated the spatiotemporal change in single-cell rheology of ascidian embryos in the animal hemisphere in the cleavage stages, using approach and retraction force curve (6,7,24‒27) and stress-relaxation AFM (28‒33) measurements, which enables obtaining mapping images of cell rheological properties in the developmental process. The results showed that single embryonic cells exhibited increased modulus and decreased fluidity in the timing of cell division. Furthermore, the correlation between the cell modulus and fluidity collapsed onto a master curve, indicating that embryonic cells interacting with the neighboring cells follow a universality of single-cell rheology during the cell cycle that involves the interphase and mitotic phases.

## MATERIALS AND METHODS

### Ascidian samples

We used adult *Ciona intestinalis* type A (*Ciona robusta*) collected around the Maizuru Fisheries Research Station, Kyoto University, and Misaki Marine Biological Station of the University of Tokyo, Japan, through the National BioResource Project (NBRP). Eggs and sperms were obtained by dissection of the ascidian gonoducts. Unfertilized eggs were dechorionated using natural seawater (Nihon Aquarium, Tokyo, Japan) containing 1.0% sodium thioglycolate (Wako Pure Chemical Industries Ltd, Tokyo, Japan.) and 0.05% actinase E (Rikaken Pharmaceutical Co. Ltd., Nagoya, Japan). After washing the dechorionated eggs in seawater several times, they were artificially inseminated and reared on 0.1% gelatin (Wako Pure Chemical Industries Ltd.)-coated dishes in seawater at 18°C for 3 h post-fertilization. The fertilized eggs were symmetrically cleavaged until the 16-cell stage and then asymmetrically divided into embryos separating animal and vegetal hemispheres (Fig. 1*B*). Cell division occurs synchronously in the animal hemisphere, still in a small number of cells, 24 cells, and a spherical-like shape is maintained until initiation of gastrulation at the late 112-cell stage in the vegetal hemisphere.

**FIGURE 1.**
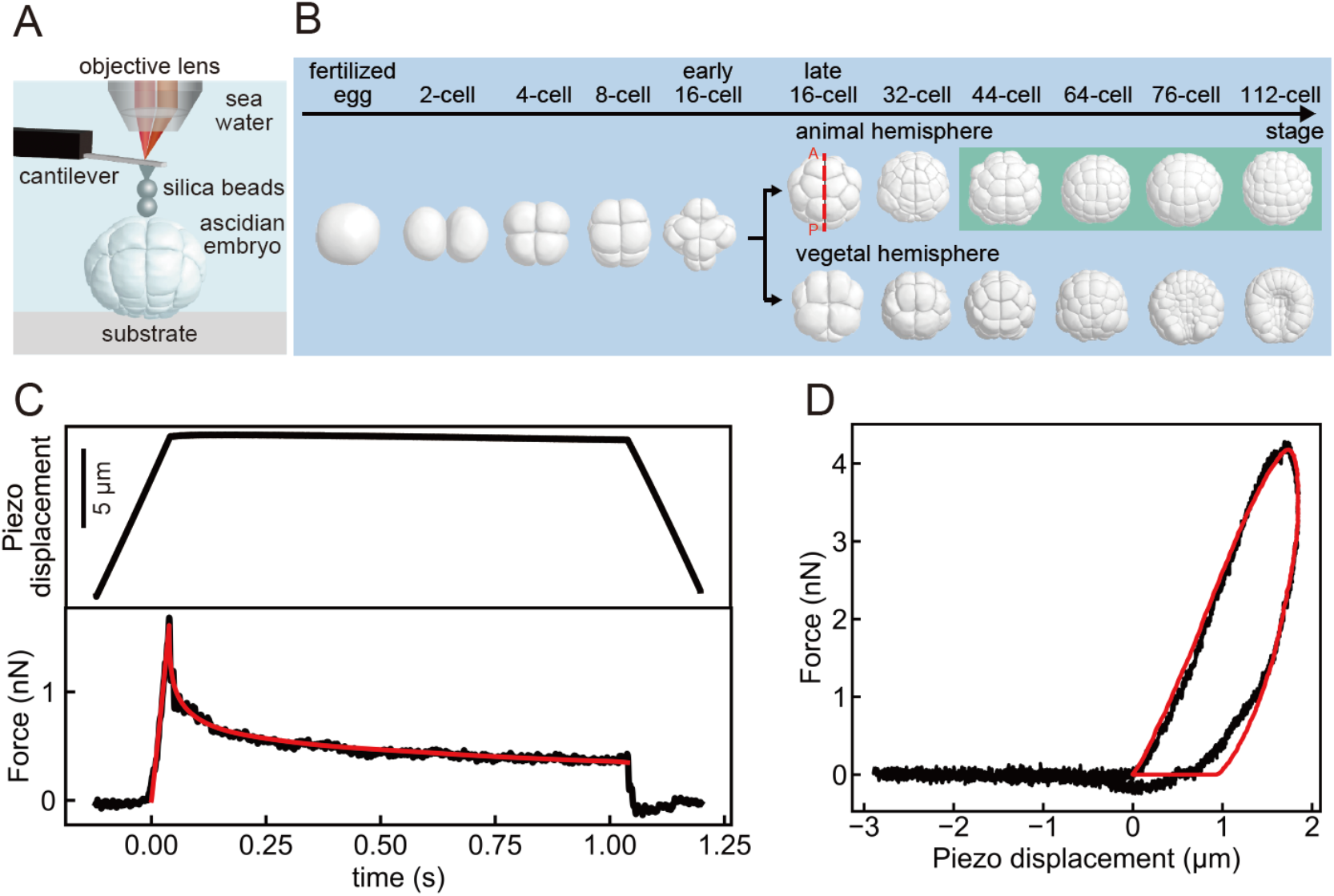
(*A*) Schematic of AFM equipped with an upright optical microscope for measuring the developing ascidian embryo weakly adhered to a dish in seawater. (*B*) Morphological changes in the 3D ascidian embryo model at early developmental stages from the fertilized egg to the 112-cell stage. Embryos from the 44-cell to 112-cell stages in the animal hemisphere (*light green*) were measured using AFM. Dotted red line represents the anterior-posterior line of the embryo. (*C*) A typical loading force and piezo displacement as a function of time during stress relaxation AFM measurement. The black dots and the red lines represent the measured data and the fitted results with Eqs. 3 and 4. (*D*) A typical approach-retraction AFM force curve. The black dots and the red lines represent the measured data and the fitted result with Eqs. 5 and 6.

### AFM measurements

A customized atomic force microscope was used to map the relative height *H* and the time-dependent relaxation modulus *E*(*t*) of ascidian embryo samples (Fig. 1*A*). The details of the AFM instrument were previously described by (22,34). Briefly, the AFM instrument was mounted on an upright optical microscope (Eclipse FN1; Nikon, Tokyo, Japan) with a liquid-immersion objective lens (plan fluor, ×10, Nikon) for the optical lever system (Nikon) and controlled with LabVIEW software (National Instruments, Austin, TX) (Fig. 1*A*). A rectangular cantilever (Biolever-mini, BL-AC40TS-C2; Olympus, Tokyo, Japan) with a nominal spring constant of <0.1 N/m was used. Before the experiments, the spring constant of the cantilever was determined in liquid environments using a thermal fluctuation procedure. To achieve a well-defined contact geometry and prevent contact between the cantilever beam and the sample surface, two silica beads with a radius *R* of ca. 2.5–3.5 μm for approach-retraction AFM and ca. 5 μm for stress-relaxation AFM were arranged tandemly from the AFM tip with epoxy glue. The AFM mapping measurements were conducted in the upper region of embryos in the animal hemisphere at 18°C in the cleavage stages from the 44-cell to 112-cell stages (Fig. 1*B*), where the embryos were weakly attached to a cell culture dish (Iwaki, Tokyo, Japan) in seawater to prevent unexpected translational movement and abnormal cell division. We obtained *H*-images corresponding to the contact points estimated from the approach force curve.

Stress-relaxation AFM and approach-retraction force curve measurements were performed to map the relaxation modulus of embryonic cells in the developmental process. In the stress relaxation experiments, the AFM cantilever probe was indented until a loading force *F*, and the force relaxation was measured while keeping indentation depth *δ* a constant (Fig. 1*C*). The scan range was approximately 72 μm × 72 μm with 4 μm spacing. The acquisition time for a single mapping image was approximately 10 min. In the approach-retraction force curve experiments (Fig. 1*D*), the scan range was approximately 75 μm × 75 μm with 3 μm spacing, and the acquisition time for a single mapping image was 2 min. To inhibit actin filament polymerization, the embryo samples were treated with 0.02 μM latrunculin-A (LatA) (Sigma-Aldrich, St Louis, MO) during time-lapse AFM mapping measurements. We analyzed the treated cells after incubation for ≥10 min with LatA to quantify the chemical dependence of cell rheology.

The cell lineages of the ascidian embryo have been determined, and the embryonic cells are invariant in space during the early developmental stages. Thus, the cells mapped using AFM were identified using the Four-Dimensional Ascidian Body Atlas database (35).

### AFM analysis

The apparent Young’s modulus *E*_Y_ of embryonic cells was estimated from approach force-distance curves using the modified Hertz contact model by taking the tilt angle *θ* of the sample surface (36). The *F* was calculated using the following formula:

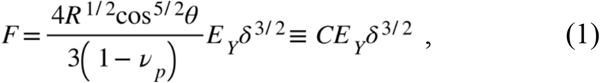

where *C* = (4*R*^1/2^ cos^5/2^*θ*/[3(1-*ν*_p_^2^]) is the factor for a spherical probe with *R*. Poisson’s ratio of the cell *ν*_p_ was assumed to be 0.5. The *θ* was estimated from the corresponding *H* image (36). The modified Hertz contact model was valid for *θ* < 40° in the case that *E*_Y_ was less than 1 kPa (36); therefore, we discussed the phenomena observed in mapping regions that satisfied *θ* < 40°.

The *E*(*t*) of a power-law rheology (PLR) model (37‒44) is expressed using the following formula:

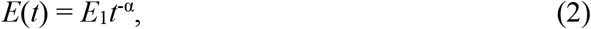

where *E*_1_ is the modulus scaling parameter at 1 s, and *α* is the power law exponent. According to the soft glassy rheology model (44,45), materials exhibited complete solid at *α* = 0 and fluid *α* = 1. Thus, *α* is referred to as the “fluidity”.

In the force relaxation experiments, the loading force as a function of time *F*(t) was analyzed with the following formula according to that used in previous studies (7,24‒ 27,42,46):

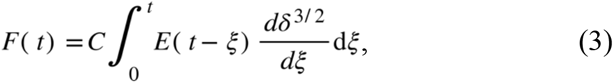

where *t* is the time initiated at the time of initial contact, and *ξ* is the dummy time variable required for the integration. The indentation depth *δ* was estimated using the following formula:

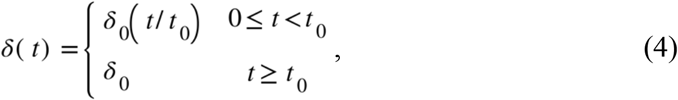

where *δ*_0_ is the maximum indentation depth at the time of *t* = *t*_0_ and *δ* is constant to be *δ*_0_ for *t* ≥ *t*_0_ (Fig. 1*C*).

We used Ting’s model to analyze the approach and retraction force curves (7,24‒27,47) using the following formula:

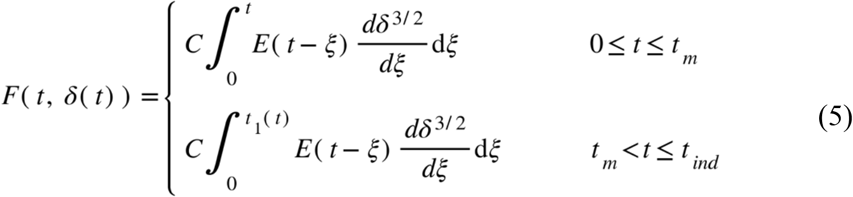

and

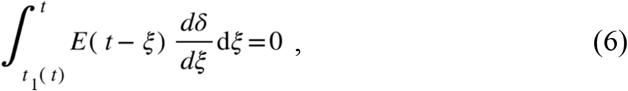

where *t*_m_ is the duration of the approach curve, *t*_ind_ is the duration of the approach and retraction curves, and *t*_1_ is the auxiliary function determined using Eq. 6 (Fig. 1*D*). *E*_Y_ and *E*_1_ images were scaled using *ν* = log_10_*E* (Pa).

## RESULTS

### Stress-relaxation AFM mapping of embryos in the cleavage stages

We first conducted stress-relaxation AFM experiments to investigate the mechanical properties of embryonic cells in the animal hemisphere from the 44-cell to 112-cell stages (Fig. 2*A*). We identified cells before and after cell division from the *H* images (Fig. 2*B*). The apparent *E*_Y_ exhibited a periodic mechanical oscillation during cell division (Fig. 2*C*), consistent with those observed in the previous study (22). Similar to *E*_Y_, the modulus scaling factor *E*_1_ significantly increased up to 200–300 Pa during cell division, then decreased to approximately 50 Pa during interphase (Fig. 2*C*). Contrastingly, the power-law exponent *α* significantly decreased to approximately 0.1 during cell division, then increased up to approximately 0.3 during interphase (Fig. 2*C*), showing that *α* has an out-of-phase response to *E*_1_. The results indicated that the change in *E*_Y_ observed in previous studies was attributed to changes in *E*_1_ and *α* rheological properties. We noticed that the stiffening and softening were not synchronized at the single cell level in single images, e.g., #2, #3, #6, and #7 in Fig. 2*B*, due to the slow mapping speed in the stress relaxation AFM.

**FIGURE 2.**
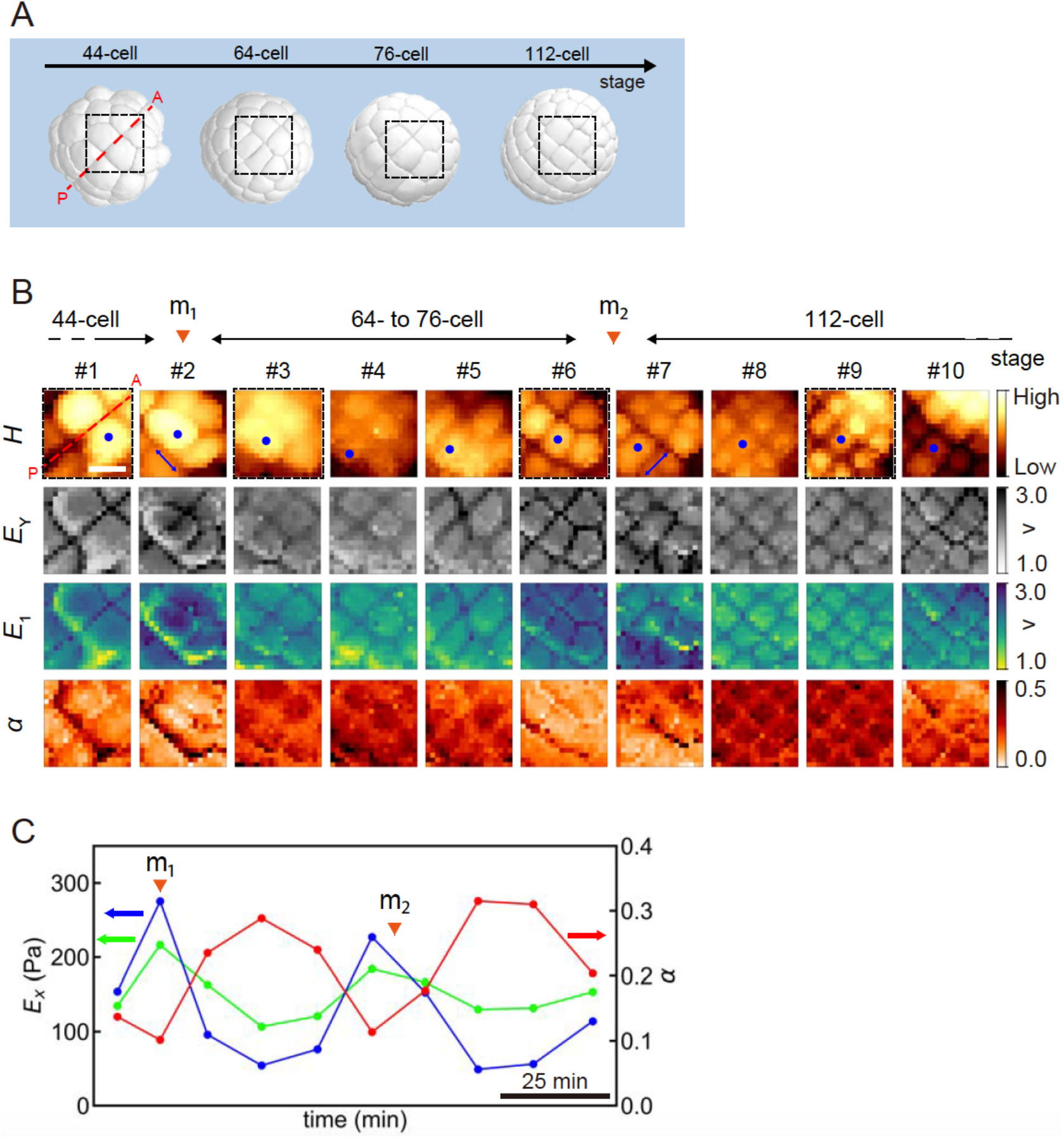
(*A*) Morphological change in animal cells of the 3D model from the 44-cell to 112-cell stages. The dotted line (*red*) represents the anterior-posterior line of the embryo. Four dotted squares (black) represent measured regions that approximately correspond to the four images (#1, #3, #6, and #9) in (*B*). (*B*) Portions of relative height *H*, apparent Young’s modulus *E*_Y_, modulus scaling factor *E*_1_, and power law exponent *α* images of the embryo mapped in the animal hemisphere from the 44-cell to 112-cell stages using stress-relaxation AFM where cells indicated by dots (*blue*) were tracked. The cell division began at times depicted by the arrowheads (*orange*: m_1_ and m_2_). Double-headed arrows (*blue*) represent the direction of cell division. Scale bar, 30 μm. (*C*) Plots of geometrically averaged *E*_Y_ (*green*) and *E*_1_ (*blue)*, and arithmetic averaged *α* (*red*) around the center of cells imaged in (*B*).

### Approach-retraction force curve mapping of embryos in the cleavage stages

Next, we measured the approach-retraction force curves with hysteresis, using Ting’s model, to investigate the rheological properties of embryonic cells in the animal hemisphere from the 44-cell to 112-cell stages (Fig. 3*A*). The mapping images exhibited a subcellular resolution, and the cell shapes in *E*_1_ and *α* images were almost identical (Fig. 3*B*). The cell locations were also consistent with those of the 3D virtual embryo model determined from confocal microscopic images (Fig. 3*A*). The results indicated that *E*_1_ and *α* analyzed from AFM data mainly reflected the rheological properties of cells that directly contacted the AFM probe. The temporal change in the rheological parameters was almost synchronized among epidermal cells during cell division (Fig. 3*B*) since the mapping images were obtained at a temporal resolution of ca. 2 min, which was higher than 10 min in the stress relaxation AFM. The correlation between *E*_1_ and *α* exhibited an out-of-phase oscillation around the mitotic phase, as observed in the stress-relaxation AFM (Fig. 2*C*). The temporal variation of rheological parameters estimated from the approach-retraction force curves varied approximately between 50–300 Pa for *E*_1_ and 0.15–0.32 for *α* (Fig. 3*C*), which was in the range of those measured by the stress relaxation AFM (Fig. 2*C*).

**FIGURE 3.**
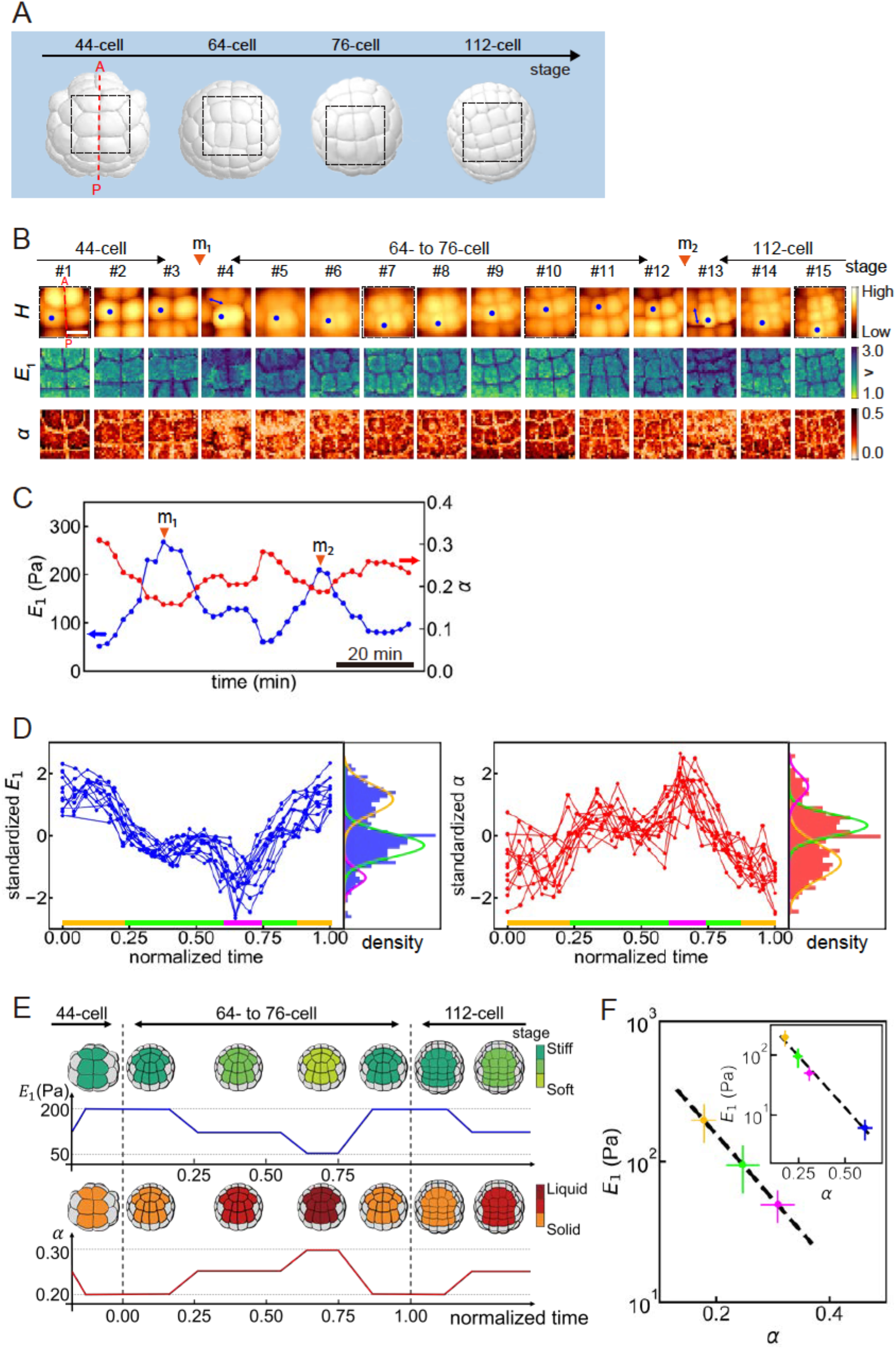
(*A*) Morphological change in animal cells of the 3D model from the 44-cell to the 112-cell stages. The dotted line (*red*) represents the anterior-posterior line of the embryo. Four dotted squares (*black*) represent measured regions that approximately correspond to the four images (#1, #7, #10, and #15) in (*B*). (*B*) Portions of *H, E*_1_, and *α* images mapped in embryo in the animal hemisphere from the 44-cell to the 112-cell stages using approach-retraction AFM force curve measurement where cells indicated by dots (*blue*) were tracked. The cell division began at times depicted by the arrowheads (*orange*: m_1_ and m_2_). Double-headed arrows (*blue*) represent the direction of cell division. Scale bar, 30 μm. (*C*) Plots of geometrically averaged *E*_1_ (*blue*) and arithmetic averaged *α* (*red*) around the center of cells imaged in (*B*). (*D*) Geometrically averaged *E*_1_ and arithmetic averaged *α* (*left*) around the center of cells were standardized for each embryo (*n* = 4). The standardized *E*_1_ (*right*) and *α* (*left*) values were plotted as a function of the normalized time between m_1_ and m_2_ in each of the cells (*n* = 14 in four embryos). On the right of each figure, the normalized distribution of the rheological values was fitted to three normal distribution functions with solid curves (*yellow, green*, and *magenta*), indicating different rheological states. The color bars represent the rheological states where the colors correspond to those of the normal distributions. (*E*) Schematic diagrams of the changes in *E*_1_ and *α* of ascidian embryonic cells in the animal hemisphere from the 44-cell to 112-cell stages. The *E*_1_ and *α* values of embryonic cells in the 3D model measured using AFM were colored without any apparent cell-to-cell heterogeneities. Cells (*light gray*) in the 3D model represent those not probed using AFM. (*F*) Plots of geometric *E*_1_ and arithmetic *α* means in rheological states for embryos (*n* = 4). The colors (*yellow, green*, and *magenta*) correspond to those in different mechanical states presented in (*D*). In the inset, the blue dot represents geometric *E*_1_ and arithmetic *α* mean measured in the LatA treatment (*n* = 3). The error bar represents the standard deviation.

### Temporal changes in power-law rheological parameters of embryonic cells during the cell cycle

To understand the detailed rheological behaviors of embryonic cells during the stages between two mitotic phases (m_1_ and m_2_), we superimposed the time evolution of rheological properties measured in different embryo samples by standardizing geometrically averaged *E*_1_ and arithmetically averaged *α* in each embryo and normalizing the time between m_1_ and m_2_ (Fig. 3*D*). We observed that in the first state of normalized time < ∼0.25 after cell division, the standardized *E*_1_ rapidly decreased while the standardized *α* increased. In the second state, between 0.25 and ∼0.6 of normalized time, the rheological parameters exhibited a plateau state. Furthermore, in the third state, between 0.6 and ∼0.75 of normalized time, the standardized *E*_1_ decreased while the standardized *α* increased. Finally, the standardized *E*_1_ increased while the standardized *α* decreased towards the cell division of m_2_. The rheological states were discriminated in three probability density functions of the PLR parameters in the time range between m_1_ and m_2_ (Fig. 3*D*), indicating that the PLR parameters of embryonic cells change in the developmental process, even during interphase. The time evolution of PLR parameters observed in the animal hemisphere (Figs. 3*B*–*D*) is summarized schematically in Fig. 3*E*. To determine the correlation between PLR parameters of embryonic cells during cell cycles, we plotted geometric *E*_1_ and arithmetic *α* means for embryos in three rheological states as presented in Fig. 3*D* (Fig. 3*F*). We found that the means *E*_1_ and *α* were plotted along a line in a semi-log plot. Moreover, the PLR parameters of embryos treated with LatA were also well located on the line (the inset of Fig. 3*F*).

## DISCUSSION

The accumulation and remodeling of actin filaments in the cortical region increase the cell stiffness beneath the cell membrane (3‒5). Microrheology experiments using optical tweezers revealed that intracellular cytoplasm softens and dissipative timescale increases during cell division (1), inconsistent with the result of this study, where embryonic cells increased the modulus and decreased viscoelastic fluidity (Fig. 3). We also observed that treatment with an actin filament polymerization inhibitor significantly decreased *E*_1_ and increased *α*. Combined, we concluded that AFM probed the rheological properties of cortical actomyosin structures rather than the interior of the cell, suggesting that the cortical actin filaments were well organized and mechanically stabilized during cell division during the developmental stages.

The rheological properties of embryos play crucial roles in developmental processes (17‒21). However, the rheology of embryo in the developmental processes has not been assessed at the single-cell level. In this study, we found that the *E*_1_ and α means of individual embryos were highly correlated (Fig. 3*F*). We also found that the PLR parameters of embryos treated with LatA collapsed into the master curve. The findings indicated that the spatiotemporal changes in the rheological properties of cells in developing embryos during the early cleavage stages were similar to those in isolated single cells *in vitro* (38‒44,48‒50).

Previous studies revealed a trajectory of mechanical parameters of isolated single cells in the cell cycle (10,51,52). AFM mechanical mapping using a melanoma cell line with a fluorescent cell cycle sensor showed that viscoelastic material properties significantly change with increasing stiffness and decreasing fluidity from the phase after cell division to the subsequent cell division (10). A real-time deformability cytometry using non-adherent leukemia cells showed that the deformability of cells increased during interphase and decreased during the mitotic phase (51). It has been reported that, in the early ascidian development stage, the cell division and morphological changes in single cells are correlated with the area of individual cell contacts through intimated communications between neighboring cells (53). Taken together, we concluded that the rheological properties of single cells under intimate cell-to-cell communications like ascidian embryos conserved the negative correlation between the cell stiffness and fluidity during the cell cycle.

Brillouin microscopy is advantageous for measuring the intracellular mechanical properties of developing embryos at the single-cell level (54‒56). Also, fluorescence microscopy observation of cell membranes in embryos is advantageous for predicting relative cell surface tensions and pressures (57). Contrastingly, our AFM allowed us to map the rheological properties of embryonic cells beneath the cell membrane as a compliant material in the developmental process, thus providing a complementary technique to unveil the whole mechanics of the embryo.

## CONCLUSION

AFM microrheology experiments such as stress-relaxation AFM and approach-retraction force curve AFM were conducted to characterize the rheological properties of ascidian embryonic cells in the animal hemisphere in the cleavage stages. The results revealed that single cells in developing embryos followed a single power-law rheology: the modulus scaling parameter, *E*_1_ increased while the fluidity *α* decreased in the timing of cell division, indicating that the cortical actin filaments were well organized and stabilized during cell division in the developmental stages. Furthermore, we found three rheological states in developing embryos during the cell cycle. The correlation between *E*_1_ and *α* during the cell cycle in embryogenesis was collapsed onto a master curve with a negative correlation, suggesting that single-cell rheology in embryogenesis conserved a universality of power-law rheology observed in single cells *in vitro*. AFM is a common technique for exploring spatiotemporal changes in the rheological properties of single cells as a compliant material in embryogenesis at the subcellular resolution.

## AUTHOR CONTRIBUTIONS

T.O. conceived and designed the experiments. T.K., T.M., M.Y., Y.T., Y.M., and Y.F. performed the experiments and analyzed the data. T.K. and T.O. wrote the paper. K.K.-S. and T.O. edited the paper.

## ACKNOWLEDGMENTS

We thank the National BioResource Project (NBRP, Japan) for *C. intestinalis* samples. We also thank Dr. Kohji Hotta (Keio University) for advice on the sample preparation. The study was supported by the JST CREST Grant Number JPMJCR22L5 and JSPS KAKENHI Grant Numbers 23K17872 and 24H00412, Japan.

## DECLARATION OF INTERESTS

The authors declare no competing interests.

## Notes

### Competing Interest Statement

The authors have declared no competing interest.

